# Preserving endothelial integrity in human saphenous veins during preparation for coronary bypass surgery

**DOI:** 10.1101/2023.08.25.554690

**Authors:** Meghan W. Sedovy, Xinyan Leng, Farwah Iqbal, Mark C. Renton, Melissa Leaf, Kailynn Roberts, Arya Malek, W. Scott Arnold, David A. Wyatt, Cynthia W. Choate, Joseph F. Rowe, Joseph W. Baker, Scott R. Johnstone, Mark Joseph

**Author notes:** **Correspondence**: Dr. Mark Joseph. Carilion Cardiothoracic Surgery 2001 Crystal Spring Ave. Suite 201, Roanoke, VA 24019. *Dr. Scott R. Johnstone. Fralin Biomedical Research Institute at Virginia Tech Carilion, 4 Riverside Circle, Roanoke, VA 24016. **Disclosure statement:** Dr. Mark Joseph is a consultant for AngioDynamics and Artivion. The remaining authors have no conflicts of interest to declare. **Informed Consent Statement:** The study protocol was approved by the local ethics committee.

## Abstract

**Objective:** While multiple factors influence coronary artery bypass graft success rates, preserving saphenous vein endothelium during surgery may improve patency. Standard methods of saphenous vein graft preparation in heparinized saline (saline) solutions result in endothelial loss and damage. Here we investigated the impact of preparing saphenous graft vessels in heparinized patient blood (blood) vs saline on cellular health and survival.

**Methods:** Saphenous vein tissues from a total of 23 patients undergoing coronary artery bypass graft surgery were split into 2 treatment groups, 1) standard preparation using saline and 2) preparation using blood. Immediately following surgery, excess tissue was fixed for analysis. Level of endothelial coverage, oxidative stress marker 4-hydroxynonenal (4HNE), and oxidative stress protective marker nuclear factor erythroid 2-related factor 2 (NRF2) expression were evaluated.

**Results:** In saline patient veins, histological analysis revealed a broken or absent luminal layer, suggesting a loss of endothelial cell (EC) coverage. Luminal cell coverage was notably preserved in blood-treated samples. Immunofluorescent staining of endothelial markers vascular endothelial cadherin (VE-cadherin) and endothelial nitric oxide (eNOS) identified a significant improvement in endothelial coverage in the blood group compared to saline. Although in both treatment groups EC expressed 4HNE indicating a similar level of oxidative stress, EC stored in blood solutions expressed higher levels of the protective transcription NRF2.

**Conclusions:** Our data indicate that maintaining and preparing saphenous vein tissues in solutions containing heparinized blood helps preserve the endothelium and promotes vein graft health. This has the potential to improve long-term outcomes in patients.

**Central Message:** During coronary artery bypass grafting, preparation of saphenous veins with heparinized saline damages the endothelium and increases oxidative stress. Heparinized blood preparation limits this endothelial loss and damage.

**Perspectives Statement:** Saphenous vein grafts are prone to failure through neointimal hyperplasia or thrombosis. Endothelial damage and loss are thought to be major contributing factors to graft failure. Here we find that preparation and preservation of saphenous vein grafts with patients’ own heparinized blood is sufficient to ensure endothelial preservation and protect vessels from oxidative stress compared with heparinized saline. These changes may increase long-term graft patency rates.

## Introduction

Neointimal hyperplasia occurs when vascular smooth muscle cells proliferate and migrate towards the vessel lumen occluding blood flow. Neointima formation is commonly found in blood vessels that have undergone surgical intervention for atherosclerosis including stent placement and coronary artery bypass graft (CABG) ^1–3^. Despite functional and morphological differences between veins and arteries, saphenous vein grafts (SVG) remain a commonly utilized conduit for complex CABG surgeries when large amounts of grafting materials are necessary. The use of SVG is associated with 3%-12% failure before discharge, 8%-25% at 1 year, and 50%-60% failure after 10 years ^4, 5^. Although this high failure rate can be partially attributed to neointima formation, SVG are also vulnerable to thrombosis which accounts for many early graft failures ^6, 7^. The risk of graft failure due to neointima or thrombosis can depend on factors such as technical handling of the vessel, inherent vessel deficiencies such as size mismatch or pathology, or patient extrinsic factors like hypercoagulability ^6, 7^. One major consideration is the combination of vein distension, cannulation, and maintenance outside of the body prior to grafting which can result in endothelial cell (EC) and smooth muscle cell (SMC) loss or damage ^8, 9^. The endothelial layer, surrounded by the glycocalyx, forms a critical barrier between circulating cells and the elastin and collagen-rich internal elastic lamina which provides structure and support in conduit vessels. Disruption of these layers can expose the elastic lamina, allowing for platelet aggregation and thrombosis ^10^. The presence of vascular EC is essential for the release of factors such as nitric oxide that prevent coagulation and thrombosis ^10^. Additionally EC damage is thought to play a role in the formation of neointimal hyperplasia ^11^. Neointima formation resulting from activated and proliferative SMC can occur as early as one-month post-operation and is exacerbated by platelet and immune cell adherence ^12, 13^.

Multiple preoperative and perioperative strategies are under investigation to improve SVG patency. Isolation strategies for harvesting SVG have been tested in randomized clinical trials including the no-touch technique versus conventional preparation methods ^7, 14^. The no-touch protocol, in which fat and surrounding tissue is not cleared from the vessel, aims to preserve the vasa vasorum and nerves in the adventitia to reduce vascular spasm, neointima, and atherosclerosis^15^. Mixed results have been documented in clinical trials, with either improved SVG patency using the no-touch technique or no significant differences, suggesting the need for additional methods of maintaining graft health ^7, 15–18^. Other notable strategies include the optimization of SVG preservation solutions. Before grafting, saphenous veins are harvested, prepared and stored outside of the body using various solutions. During this time, nutrient deprivation can lead to EC dysfunction and loss ^19–22^. As outlined above, the various downstream risks of SVG thrombosis, neointima formation, and atherosclerosis are rooted in this EC injury. It is thought that methods to maintain the EC layer during CABG can improve SVG patency and longevity.

Heparin-supplemented saline solution is commonly used for SVG preservation over an extended period of time during surgical preparation. However, there is documented evidence for detrimental effects of saline on endothelial cell function ^23^. Despite this, many institutions still use saline as their primary SVG preparation and maintenance solution ^23^. Autologous arterial heparinized blood (blood) as an alternative preparation solution is potentially favorable due to the close replication of an *in vivo* microenvironment. Despite the physiological relevance, reports of enhanced endothelial preservation are conflicting ^24, 25^. Studies have also characterized endothelial cell morphology using electron microscopy, graft wall tension, oxidative stress, and apoptosis, in saline vs blood stored saphenous veins ^26–28^. Although they saw some benefit to endothelial preservation in blood, these studies did not assess levels of endothelial coverage and more research is required to understand the underlying causes of EC death ^26–28^.

In this report, histological and immunofluorescent analysis reveals preservation of both the luminal EC layer and the improved cellular coverage of internal elastic lamina of SVG stored in blood versus saline for CABG. Our data also demonstrate a potential protective effect in blood-preserved veins, with increases in antioxidant effects limiting vulnerability to oxidative stress. Because vessels were fixed at the conclusion of each surgery, we believe this analysis provides a more accurate view of the typical saphenous vein at the time of grafting. Ultimately, our results provide evidence to support the use of arterial blood for graft storage during surgery and for the standardization of graft preservation protocols during CABG procedures.

## Methods

### Surgical coronary artery bypass procedure

Standard techniques were used in the harvest of all saphenous veins. All veins were harvested by surgical first assists or physician associates that were experienced, using endoscopic instruments (Hemopro, Maquet) and techniques. Once veins were harvested, heparinized saline or heparinized arterial blood was used to test the vein quality and for preservation. All veins were cannulated and used in a reversed fashion. For patients who underwent preservation using heparinized saline, a standard mixture of 10K units of heparin mixed with 1 liter of normal saline was used. For patients who underwent use of heparinized blood, a total of 100cc of heparinized arterial blood was used for injection and preservation. Any unused segment of vein was preserved in the solution it was prepared in until the conclusion of the procedure. At the conclusion of the surgery, all remaining veins were immediately fixed in 10% formalin.

### Consent and Inclusion Procedure

The Institutional Review Board (IRB) at the Carilion Clinic approved this procedure in Dec of 2020.

### Histological analysis

Fixed, SVG samples were processed for immunohistochemistry and immunofluorescent staining using standard paraffin embedding protocols. For H&E and Movats Pentachrome staining, SVG tissue sections were deparaffinized and stained using standard procedures.

### Immunofluorescence

To detect the presence of EC, SVG tissue sections were blocked in a phosphate buffered saline solution containing 0.1% NP-40 and 1% BSA followed by overnight incubation in antibodies targeting human VE-cadherin (Catalog No. MABT886, 1:100 dilution, Sigma) or endothelial nitric oxide synthase (eNOS, Catalog No. 610296, 1:100 dilution, BD Biosciences). To detect oxidative stress and antioxidant effects, samples were incubated with antibodies targeting 4-hydroxynonenal (4HNE, ab48506, 1:100, Abcam) and nuclear factor erythroid 2-related factor 2 (NRF2, #12721, 1:25, Cell Signaling Technology). Antibody binding was identified by incubation with goat anti-rabbit or anti-mouse secondary antibodies (Thermo Fisher Scientific, 1:2,000, A-21237 and A-21246) as well as DAPI to counterstain nuclei. Secondary antibody controls for each sample were used to confirm antibody specificity and background fluorescence levels. Samples were mounted using FadeStop^TM^ fluorescent mounting reagent with DAPI (GenuIN Biotech #272).

### Imaging and analysis

Histological imaging was completed on a Nikon Ci-L histology microscope using tile scan functions to capture whole tissues. Immunofluorescence was imaged on a Nikon NIE confocal microscope. Maximum intensity Z-stack projection along with tile scanning was used to visualize entire samples in one image. All quantitative analysis was performed using NIS Elements software (Nikon). Data for quantification of EC coverage was performed by a researcher blinded to sample identification.

EC coverage was measured as a proportion of coverage by VE-cadherin and eNOS positivity. The proportion of EC coverage was calculated by measuring the length of the entire vessel lumen as well as the length of eNOS/VE-cadherin positive luminal areas. Positive/negative areas were determined visually, with secondary controls used to define background signal.

To determine levels of 4HNE and NRF2, EC cell borders (where present) were detected using NIS Elements thresholding capabilities. Ten detected cells were selected for measurement per sample, and the mean 4HNE signal per cell was quantified and averaged. NRF2 measures were performed in a similar manner except that thresholding to DAPI was used to detect the borders of cell nuclei.

### Statistical analysis

Statistical analysis was performed using Graphpad Prism, Unpaired two-sample t-tests were used to assess differences between treatment groups.

## Results

### Histological analysis of saphenous vein graft tissues reveals damage in saline treated veins

A schematic of vessel cannulation, perfusion with either saline or blood, and fixation of leftover SVG tissue occurring immediately after surgery is shown in **Fig. 1A**. The luminal edge of saline and blood samples (n=6) were analyzed by hematoxylin and eosin stain (**Fig. 1B**, **Supplemental Fig. 1**) with nuclei appearing in dark purple. In vessels, H&E staining revealed a reduction in cell coverage at the luminal edge of the vessel in saline samples, with greater cell coverage observed in blood samples, suggestive of saline-induced endothelial cell loss. Movats Pentachrome stain (n=6) demonstrates increases in exposed collagen and elastin layers at the luminal edges of vessels in saline compared to blood samples (**Fig. 1C, Supplemental Fig. 2**).

**Figure 1:**
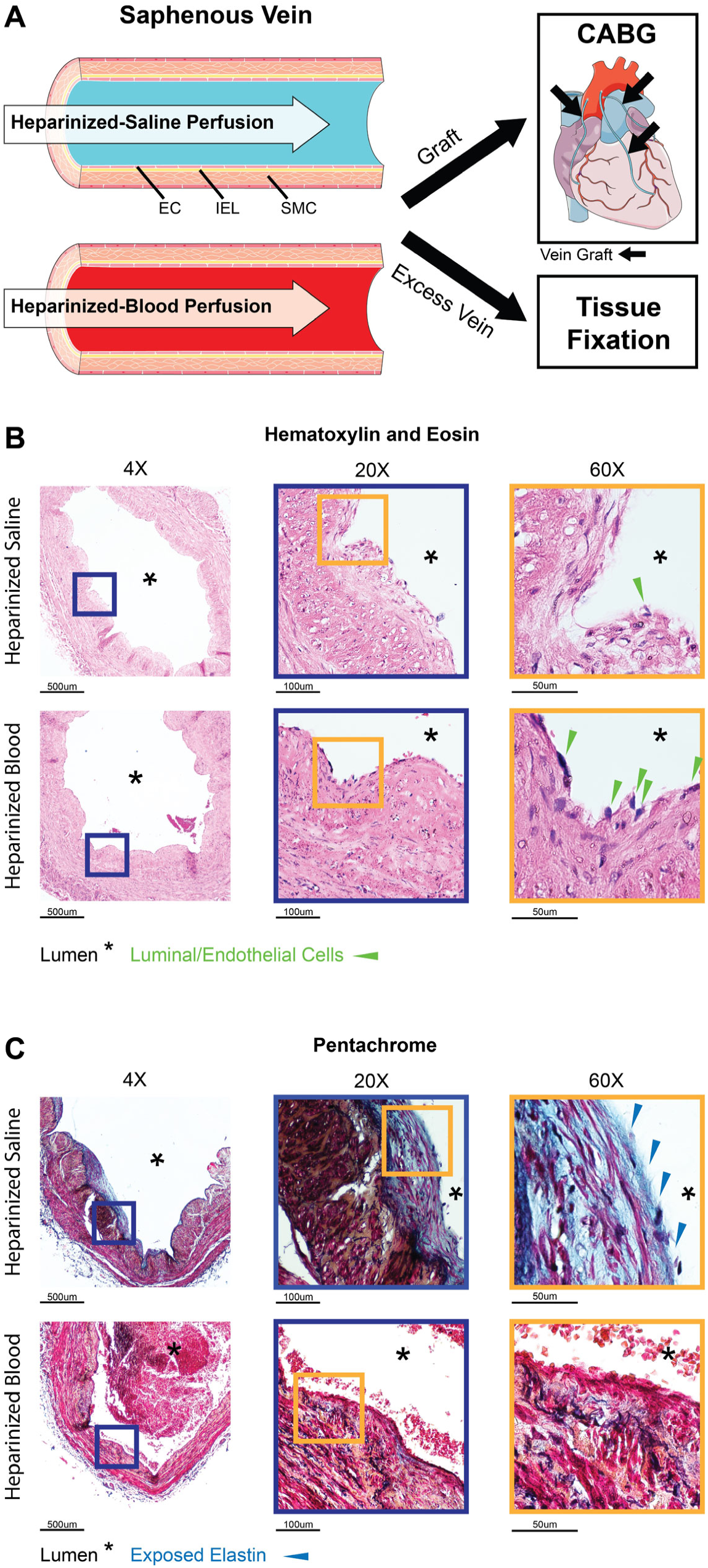
Saline exposure results in SVG damage. (A) Illustration of saphenous vein *ex vivo* maintenance, use in grafting procedures, and fixation for inclusion in the present study. Leftover SVG tissues were fixed in 10% formalin immediately following surgery. (B) Representative images of different magnifications of H&E stained saphenous veins prepared in either saline or blood. Green arrows indicate cell nuclei (purple) at the luminal edge of the tissue likely to be endothelium. (C) Representative images of different magnifications of Movats Pentachrome stained saphenous veins maintained in either saline or blood. Blue arrows represent areas of tissue that appear to be damaged with exposed elastin (blue).

### Endothelial coverage is improved using heparinized patient arterial blood

To determine if the reduction in luminal edge cells in saline samples was due to endothelial loss, SVG tissues were stained for specific endothelial cell markers VE-cadherin (n=7, **Fig. 2A, Supplemental Fig. 3**) and eNOS (n=7, **Fig. 2B, Supplemental Fig. 4**). Both measurements identified endothelial marker coverage was near 100% in blood-maintained samples while coverage in saline samples was significantly reduced (approximately 20 to 80%). Visual inspection of whole vessel sections reveals that the remaining saline-exposed endothelium appears clumped together, broken up by sections of endothelial-denuded vessels.

**Figure 2:**
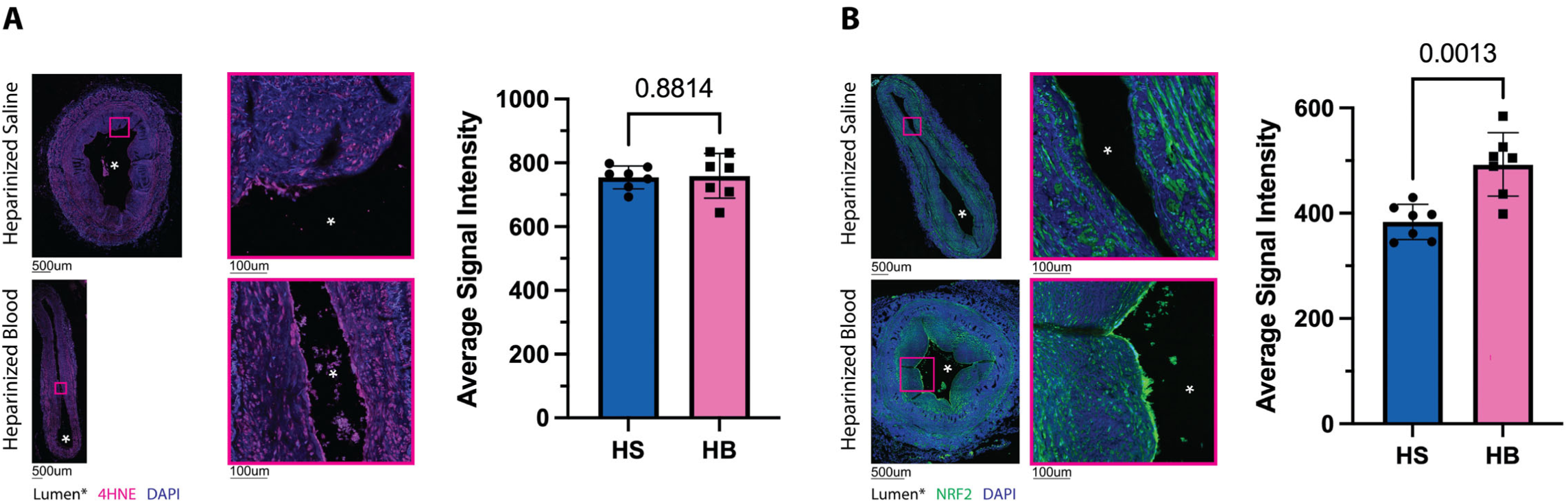
SVG preservation in blood improves vessel endothelial coverage. Representative images of immunofluorescent detection of endothelial markers (A) VE-cadherin and (B) eNOS expression (n=7). EC markers are shown in pink while DAPI labeled nuclei are shown in blue. Lower magnification images of the whole vessels contain pseudo-color lines indicating staining positive areas in orange and enlarged images contain orange arrows indicating staining positivity. Pink boxes indicate the amplified image area. Graphs represent the percentage of each vessel lumen determined to be positive for endothelial markers.

### Heparinized patient blood is protective against oxidative stress

Due to the apparent damage caused during ex-vivo vein preparation and preservation, we investigated if oxidative stress could contribute to endothelial damage. Veins were stained for the oxidative stress marker, 4HNE (**Fig. 3A, Supplemental Fig. 5**). We observed signal for 4HNE in both saline and blood samples, with no apparent differences in 4HNE signal between saline and blood treated endothelium. We then tested for the presence of the antioxidant NRF2, a transcription factor known to be protective against oxidative stress-induced damage. Our data indicate a significant elevation of NRF2 in blood-SVG endothelium only, with almost no expression detected in the remaining endothelium of saline patient samples (**Fig. 3B, Supplemental Fig. 6**).

**Figure 3:**
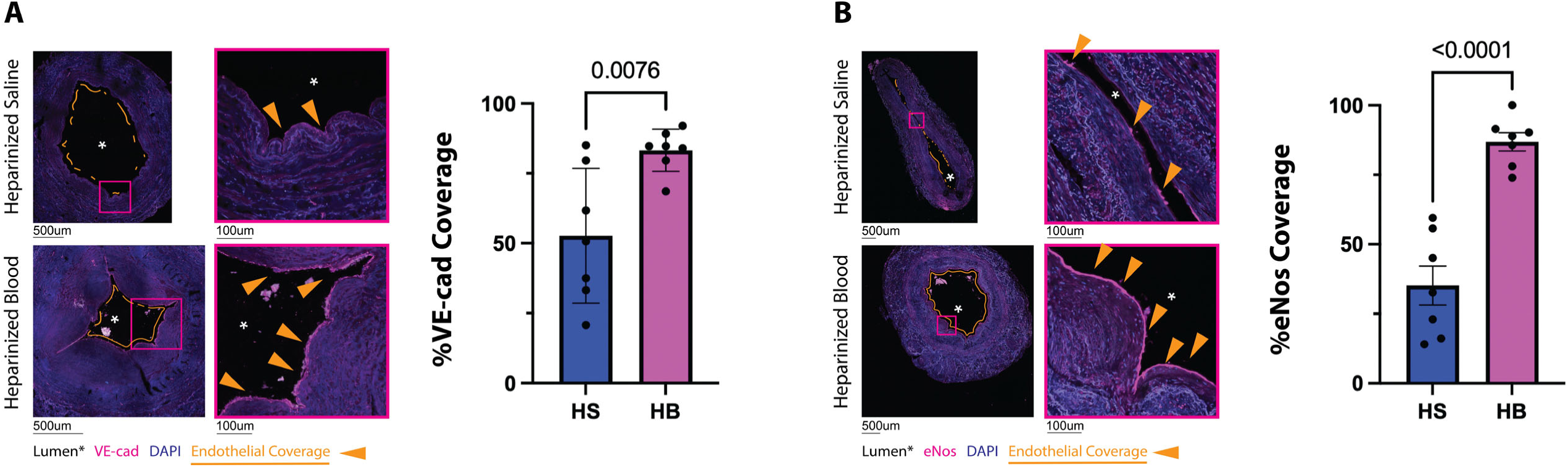
SVG preservation in blood increases resistance to oxidative stress. Representative images of immunofluorescent detection of (A) 4HNE (pink) and (B) NRF2 (green) expression (n=7). Pink boxes indicate the amplified image area. Graphs represent calculated 4HNE and NRF2 signal intensity averages in luminal EC.

## Discussion

CABG using the greater saphenous vein is often a preferred intervention for severe coronary artery disease, making SVG patency extremely important for patient health and survival. Even so, SVG long-term patency rates can be as low as around 50% ^4, 29^. Although a healthy endothelium is essential for preventing graft failure, the direct causes and mechanisms of endothelial damage remain unclear. In this study, we assess the impact of standard saline SVG preparation and preservation methods versus an inexpensive and easy-to-implement autologous arterial blood preservation method on saphenous vein endothelium. Histological analysis revealed fragmented luminal cell coverage and exposed collagen layers in veins maintained in saline, which was prevented with use of blood for preparation and storage. We confirmed this was due to a loss of continuous endothelium by immunofluorescent techniques. A key question was whether the remaining endothelium are healthy following surgery. Our data indicate that the use of blood increases antioxidative signaling in the endothelial layer, which may improve cellular recovery after use. This pattern of endothelial loss and increase in oxidative stress, exacerbated by vessel preparation and maintenance in heparin saline, may be a key factor in SVG failure rates.

The endothelium is a critical component in modulating vascular health and resisting thrombosis. Therefore, it is essential that it be preserved during surgery to maintain an effective barrier. The process of re-endothelialization is relatively slow and data from murine endothelial injury models suggests this occurs by both migration and proliferation of existing endothelial cells as opposed to circulating endothelial progenitor cells ^30^. Thus, limiting endothelium loss is critical to long-term vessel healing after surgery. Our data suggest that significant endothelial loss occurs in saline prepared SVG. This may occur for several reasons including the use of heparin and a lack of nutrients for an extended period of time. The endothelium of blood vessels is protected by a thin proteoglycan layer called the glycocalyx, essential for regulating vascular permeability, clot formation, and the response to environmental changes including nitric oxide signaling ^31^. Previous studies have demonstrated the vulnerability of this layer to heparin, which impairs glycocalyx function resulting in increased vascular leak and impaired shear induced vasodilation ^32^. Endothelial cells are also highly susceptible to their environment, with studies in culture demonstrating significant cellular dysfunction in saline exposed cultured saphenous vein EC ^23^. Thus, it is likely that the combination of dysfunctional glycocalyx and nutrition deficiency lead to cell death in the endothelium during the surgical preparation. While a majority of CABGs done routinely require the use of heparin to prevent coagulation while on cardiopulmonary bypass, our data indicate that the presence of autologous blood is beneficial in resisting endothelial loss.

While the endothelium is preserved in our blood samples, it was possible that we only limit immediate cellular death, and that the endothelium may degrade at a later point. Although it is not possible in our study to look at long term effects, we aimed to address the overall health of the endothelium at the time of use. Increases in cellular reactive oxygen species (ROS) occur when cells are unable to counteract production of ROS, a normal byproduct of cellular energy production. This can be due to cellular dysfunction induced by environmental stressors leading us to hypothesize that exposure to saline may result in endothelial susceptibility to oxidative stress^33^. Ultimately, the effect of uncontrolled ROS increases is damage to proteins, nucleic acids, and organelles that can result in apoptosis ^34^. Interestingly, our data indicate the presence of ROS (4HNE) in both saline and blood samples at similar levels. ROS-induced stress is controlled via the production of antioxidant enzymes. Therefore, we tested whether either condition altered activation of antioxidant pathways ^35^. A significant contributor to antioxidant defense is NRF2, a transcription factor that increases in response to oxidative stress and promotes the production of antioxidants ^35^. Our data demonstrate a clear and significant upregulation of NRF2 expression in the blood EC samples. These data suggest that the increase in endothelial retention in blood-exposed veins is at least partially due to an increased resilience to the damaging effects of oxidative stress, which may have lasting effects on endothelial health and wound repair following surgery. One potential limitation of this study is not being able to control for potential damage to the vein during preparation by avoiding overdistention. Although the technique for harvest and preparation was the same in both groups of patients as they had the same assistants preparing the vein, we were unable to objectively measure the pressure at which the veins were distended in either group. Ultimately this means that it is possible that individual differences in surgical technique could have influenced the results of this study. Additionally, a low sample size makes results hard to generalize on a population level. Finally, we were unable to follow up on patient outcomes after each surgery. Future studies should include long-term graft patency data to better evaluate the link between endothelial health and patient outcomes.

Although we find endothelial cells in blood maintained SVG to be more prevalent and less vulnerable to cellular stress, a number of vein grafting challenges remain. The first of these is damage sustained by smooth muscle cells. Smooth muscle is the primary contributor to the formation of neointima after vascular damage ^36^. Endothelial damage is thought to drive smooth muscle cell activation leading to neointima formation ^11^. As such, preserving endothelium using blood preparation methods may result in clinical benefit in the form of reducing smooth muscle driven neointima formation. Even so, it is impossible to rule out the contribution of damage sustained directly by smooth muscle to neointimal formation. It is also possible that blood preparation protects the smooth muscle itself from saline induced damage. This is suggested by the fact that our data indicates an increase in NRF2 expression in SMC layers of vessels, but this requires further investigation. Regardless, it remains important to minimize smooth muscle damage using already developed methods such as no-touch vessel preparation techniques. Additionally, while we show endothelial preservation during SVG procedures with blood preparation, some level of endothelial damage and loss is inevitable. As such, it is important that future research addresses the mechanisms of endothelial repair to provide optimal post-procedure treatments that prolong SVG life. We are hopeful that the findings in this study will inspire the use of improved SVG preparatory solutions such as autologous blood and ultimately translate into clinical benefit for the thousands of patients who undergo CABG procedures in the US each year^37^.

## Glossary of Abbreviations

CABG: Coronary artery bypass graft
EC: Endothelial cell
SVG: Saphenous vein grafts
4-HNE: Nuclear factor erythroid 2-related factor 2
NRF2: 4-hydroxynonenal
eNOS: Endothelial Nitric Oxide
VE-cadherin: Vascular endothelial cadherin

## Author contributions

MS performed experimentation, and data analysis and wrote the manuscript

XL performed experimentation, and data analysis and wrote the manuscript

FI performed experimentation, and data analysis and wrote the manuscript

MR performed data analysis and wrote the manuscript

ML performed experimentation, and data analysis and wrote the manuscript

KR performed data analysis and wrote the manuscript

AM performed data analysis and wrote the manuscript

WSA performed patient surgeries and wrote the manuscript

DW performed patient surgeries and wrote the manuscript

JR performed patient surgeries and wrote the manuscript

CC performed patient surgeries and wrote the manuscript

JWB performed patient surgeries, reviewed all data, and wrote the manuscript

SRJ conceived the study, reviewed all data, and wrote the manuscript

MJ conceived the study, performed patient surgeries, reviewed all data, and wrote the manuscript

## Acknowledgements

Special thanks to Kimberly Briggs, RN and the operating room staff in their assistance with this study.

---

**Supplemental Figure 1: All H&E stained SVG sections.** All H&E stained sections with whole vessels shown in 4x, 20x and 60x magnification (n=6). Blue and yellow boxes show the magnified area on the preceding image. Green boxes indicate samples used in figure 1.

**Supplemental Figure 2: All Movat Pentachrome stained SVG sections.** All Pentachrome stained sections with whole vessels shown in 4x, 20x and 60x magnification (n=6). Blue and yellow boxes show the magnified area on the preceding image. Green boxes indicate samples used in Figure 1.

**Supplemental Figure 3: All VE-cadherin stained samples.** Each image was taken on 20x magnification with VE-cadherin in pink and DAPI in blue. For each patient, the left image shows the entirety of the stained vessel with a pink box to indicate the area of the enhanced magnification to the right. Green boxes indicate images used in Figure 2.

**Supplemental Figure 4: All eNOS stained samples.** Each image was taken on 20x magnification with eNOS in pink and DAPI in blue. For each patient, the left image shows the entirety of the stained vessel with a pink box to indicate the area of the enhanced magnification to the right. Green boxes indicate images used in Figure 2.

**Supplemental Figure 5: All 4HNE stained samples.** Each image was taken on 20x magnification with 4HNE in pink and DAPI in blue. For each patient, the left image shows the entirety of the stained vessel with a pink box to indicate the area of the enhanced magnification to the right. Green boxes indicate images used in Figure 3. Staining specificity was confirmed using secondary only control staining.

**Supplemental Figure 6: All NRF2 stained samples.** Each image was taken on 20x magnification with 4HNE in green and DAPI in blue. For each patient, the left image shows the entirety of the stained vessel with a pink box to indicate the area of the enhanced magnification to the right. Green boxes indicate images used in Figure 3. Staining specificity was confirmed using secondary only control staining.

