## Supplemental Figures 1-6 for "Preserving endothelial integrity in human saphenous veins during preparation for coronary bypass surgery"

Supplemental Fig 1:

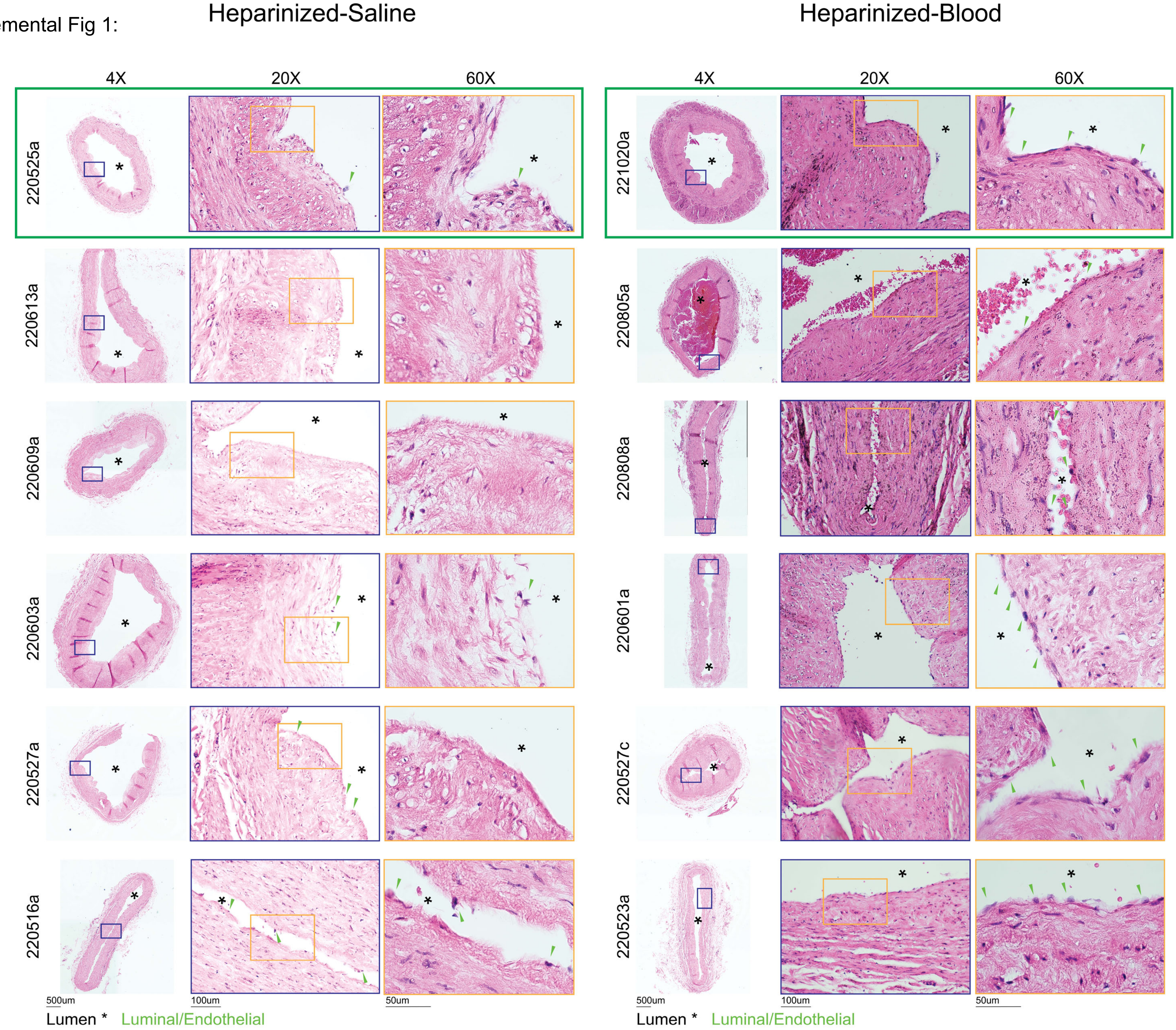

Supplemental Fig 2:

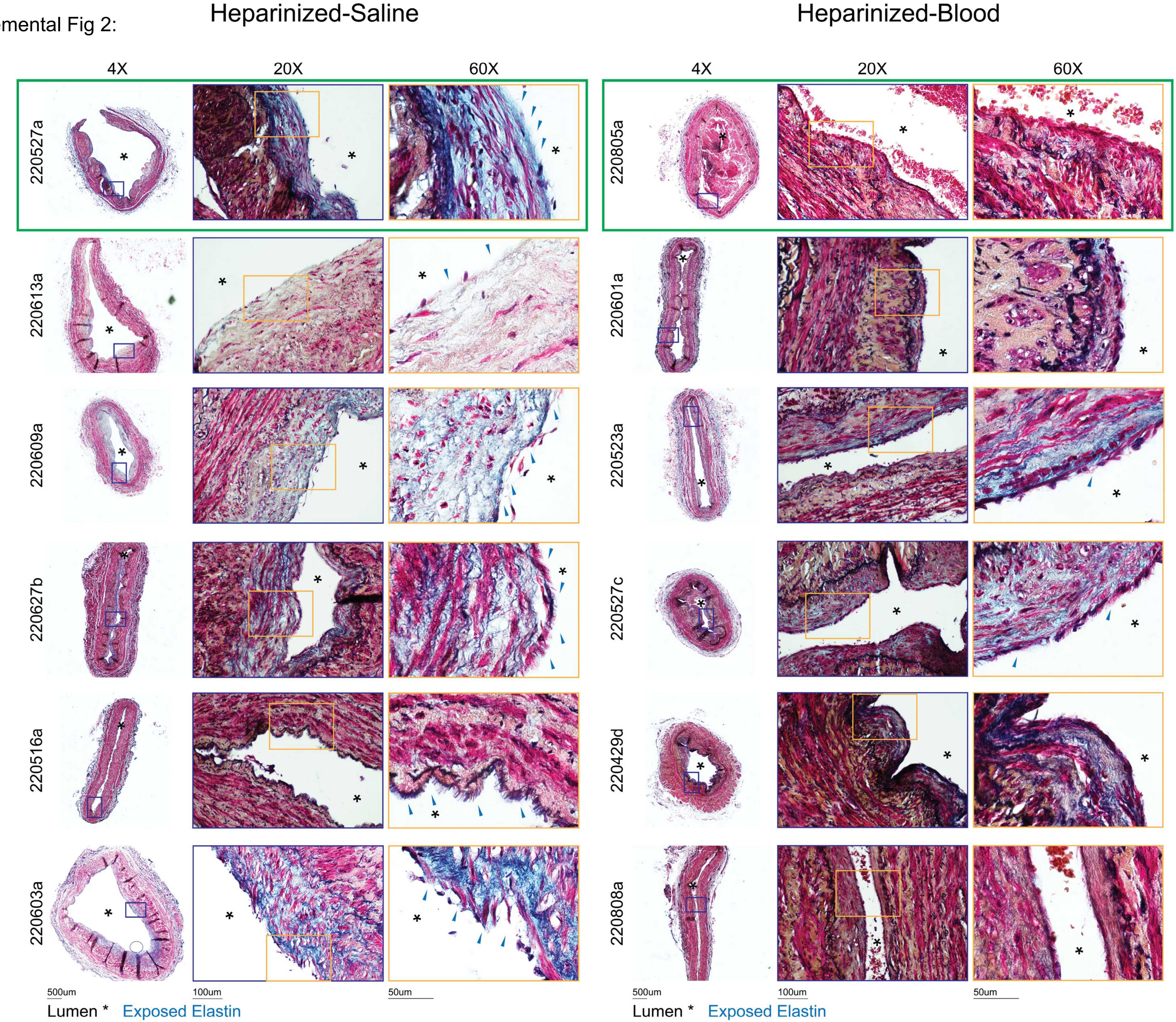

Supplemental Fig 3:

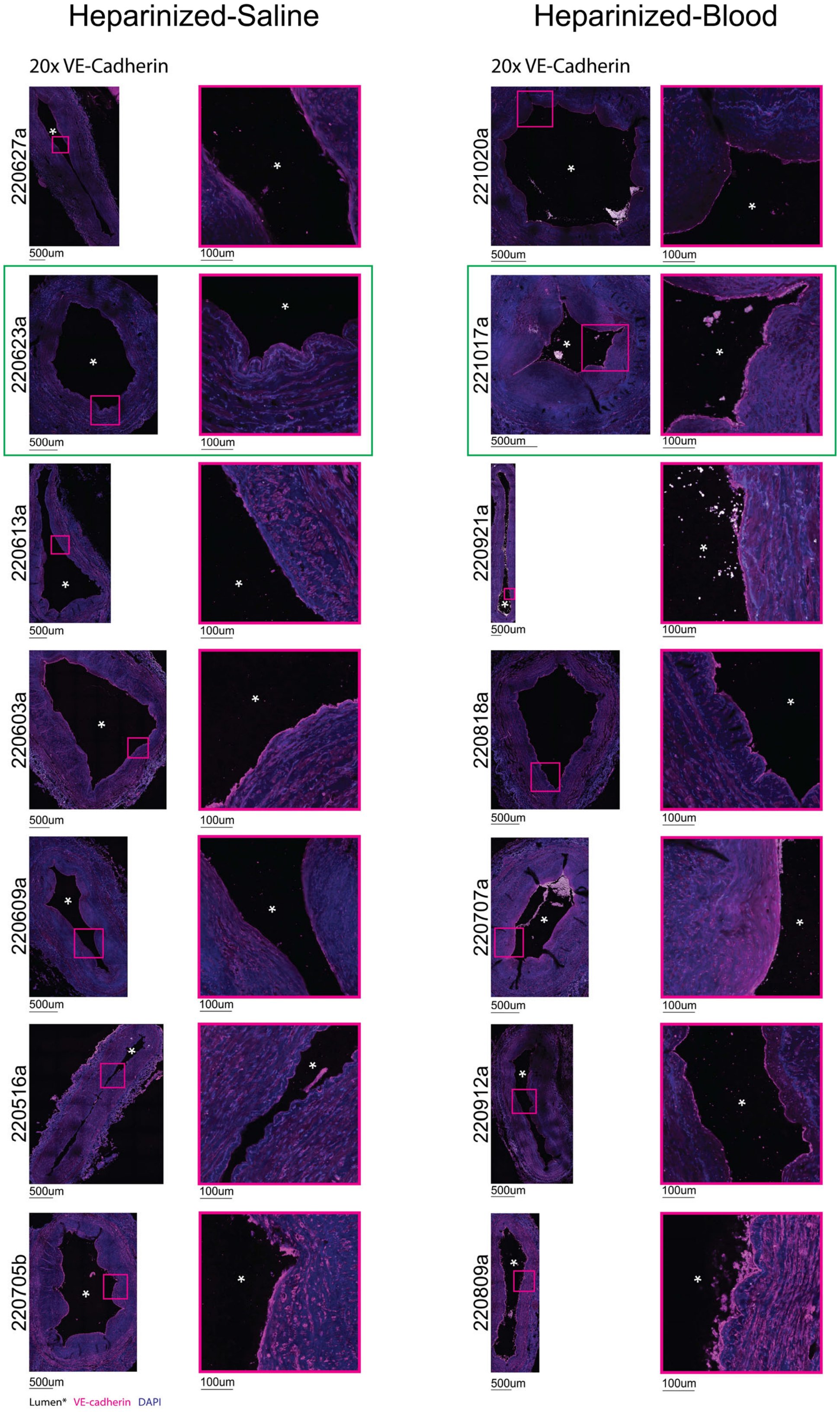

Supplemental Fig 4:

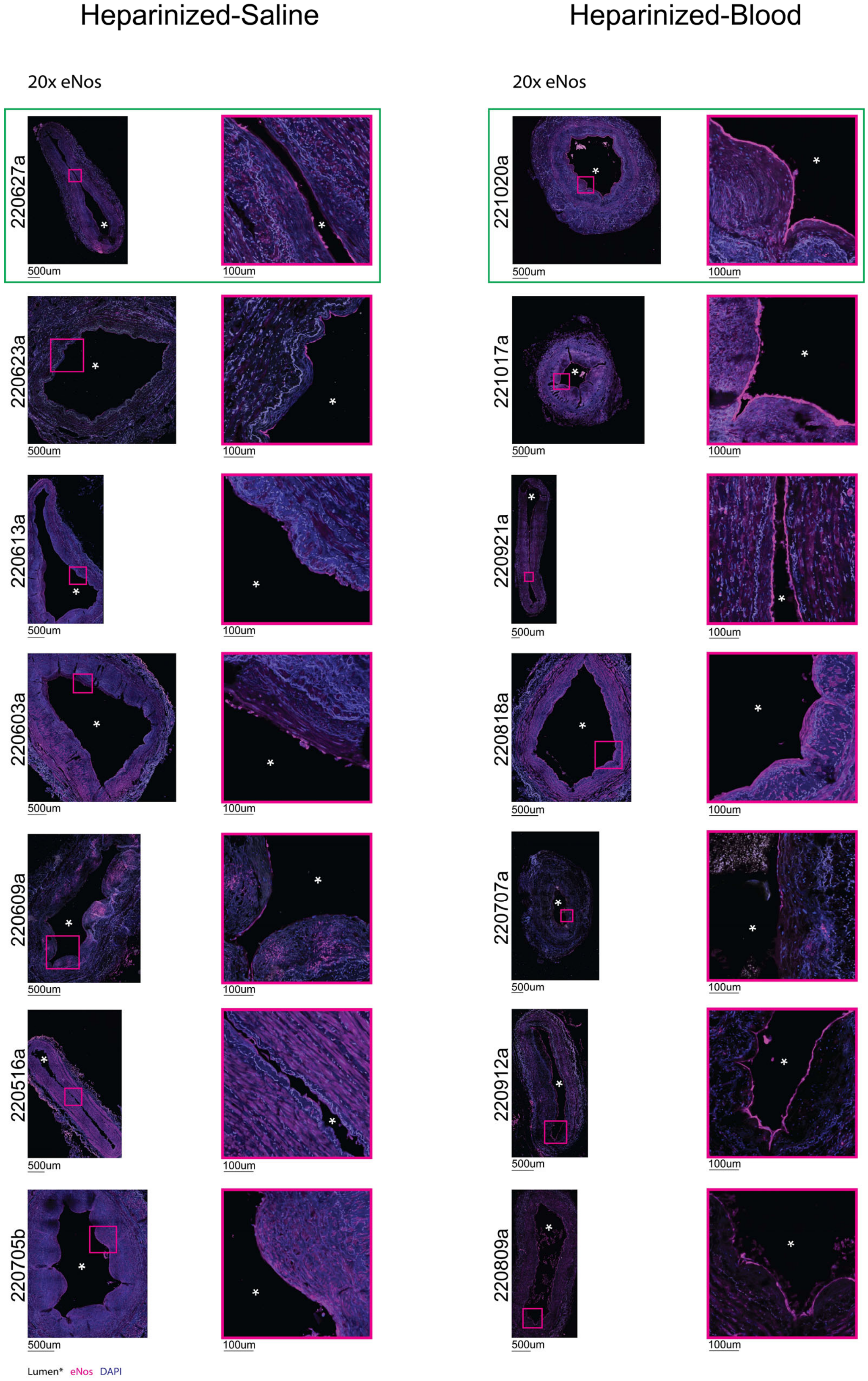

Supplemental Fig 5:

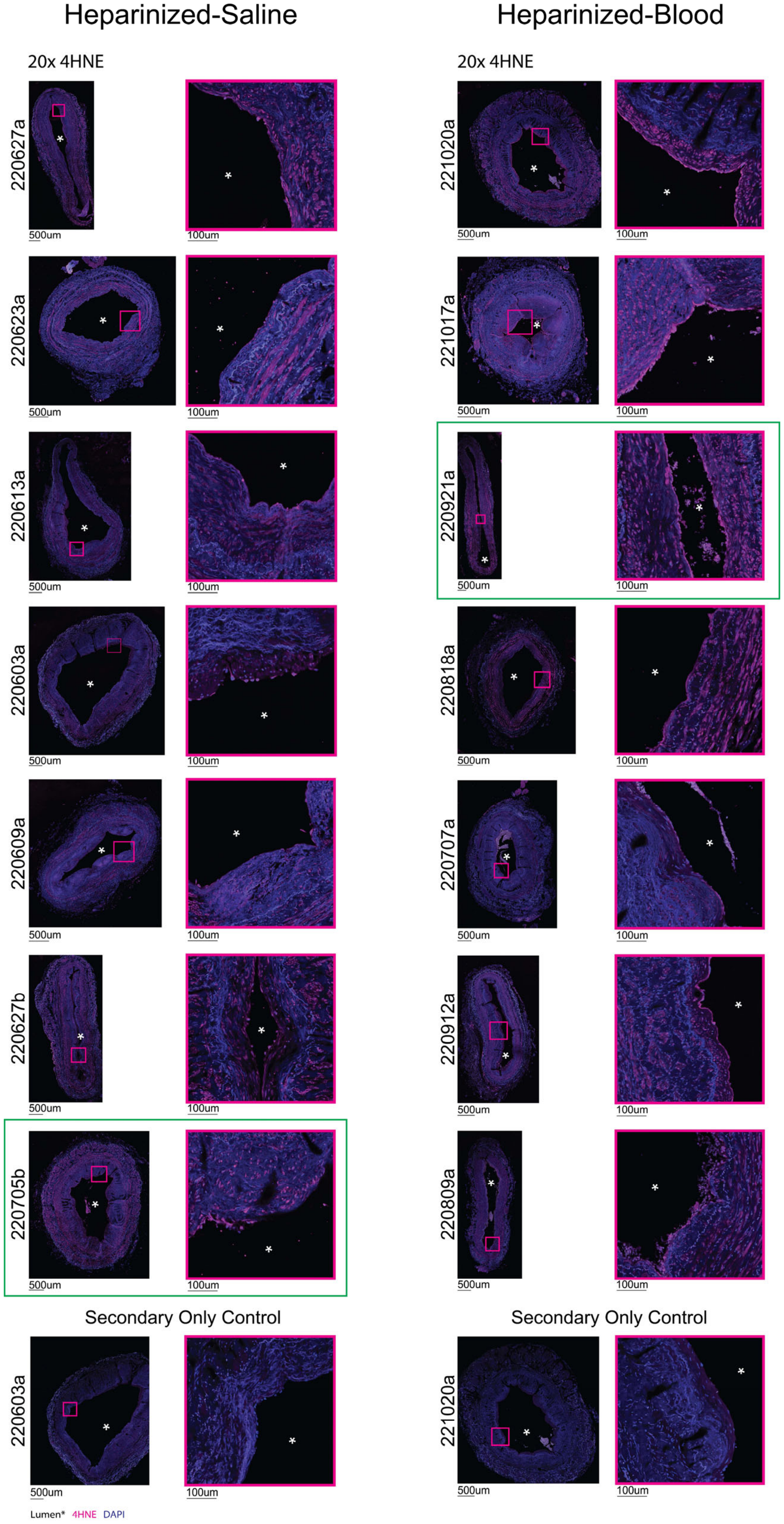

Supplemental Fig 6:

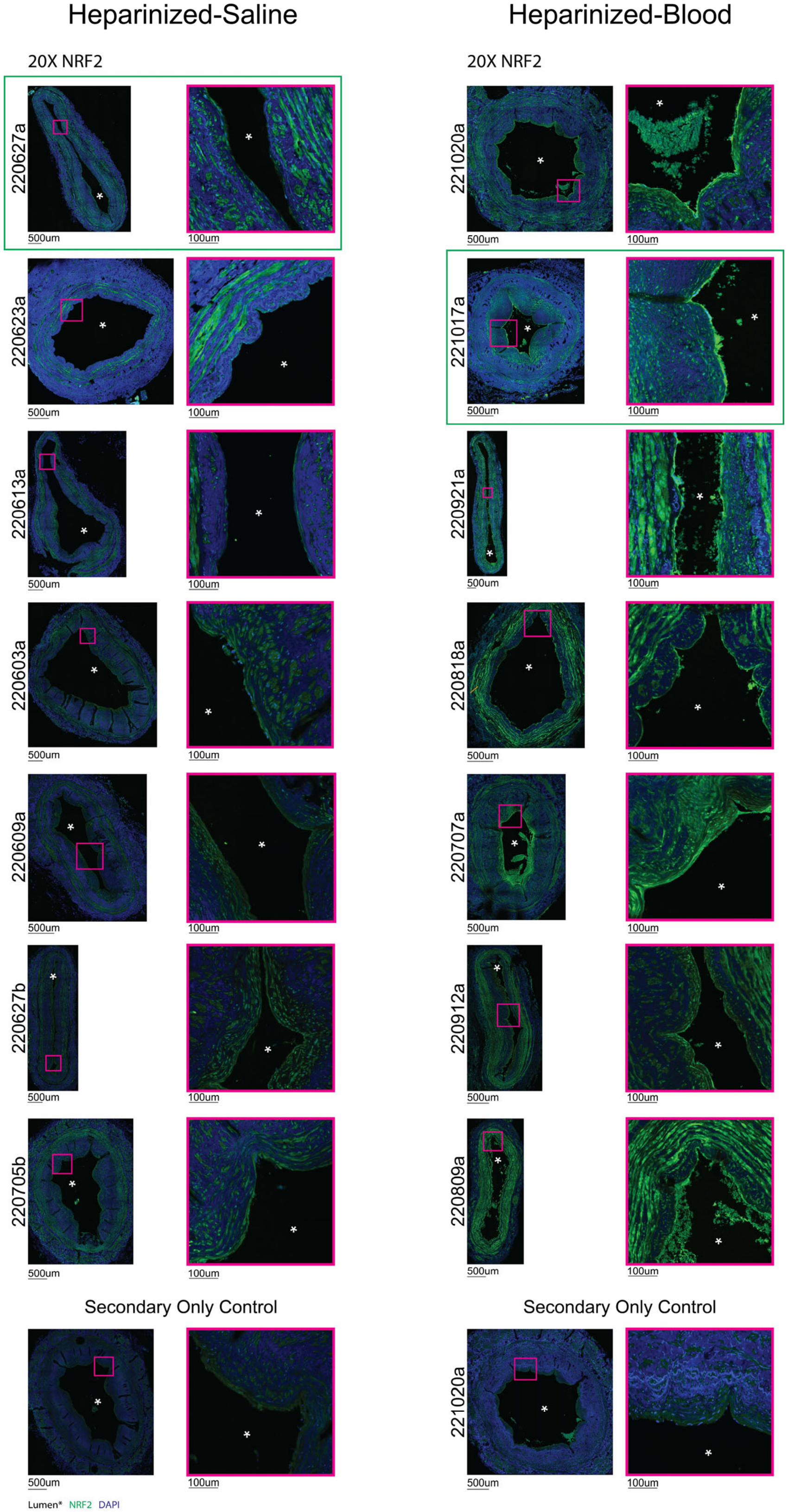
